# Functional validation of transposable element derived *cis*-regulatory elements in Atlantic salmon

**DOI:** 10.1101/2022.11.02.514921

**Authors:** Hanna M. Sahlström, Alex K. Datsomor, Øystein Monsen, Torgeir R. Hvidsten, Simen Rød Sandve

**Affiliations:** Department of Animal and Aquacultural Sciences, Faculty of Bioscience, Norwegian University of Life Sciences; Faculty of Chemistry, Biotechnology and Food Science, Norwegian University of Life Sciences, Ås, Norway

## Abstract

**Background:** Transposable elements (TEs) are hypothesized to play important roles in shaping genome evolution following whole genome duplications (WGD), including rewiring of gene regulation. In a recent analysis, duplicate gene copies that had evolved higher expression in liver following the salmonid WGD ~100 million years ago were associated with higher numbers of predicted TE-derived *cis*-regulatory elements (TE-CREs). Yet, the ability of these TE-CREs to recruit transcription factors (TFs) *in vivo* and impact gene expression remains unknown.

**Results:** Here, we evaluated the gene regulatory functions of 11 TEs using luciferase promoter reporter assays in Atlantic salmon (*Salmo salar*) primary liver cells. Canonical Tc1-*Mariner* elements from intronic regions showed no or small repressive effects on transcription. However, other TE-derived *cis*-regulatory elements upstream of transcriptional start sites increased expression significantly.

**Conclusion:** Our results question the hypothesis that TEs in the Tc1-*Mariner* superfamily, which were extremely active following WGD in salmonids, had a major impact on regulatory rewiring of gene duplicates, but highlights the potential of other TEs in post-WGD rewiring of gene regulation in the Atlantic salmon genome.

## Introduction

Whole genome duplications (WGD) is believed to result in functionally redundant genes, which can evade selective constraints and thereby evolve novel functions and regulation (1). In vertebrates, a series of recent studies across four ancient WGDs have revealed that regulatory divergence is extensive and mostly asymmetric following WGD (2–4), with one gene copy retaining an ancestral-like regulation while the other copy evolves novel regulatory phenotypes. Yet, the underlying mechanisms driving evolution of novel regulatory phenotypes are mostly unknown.

Salmonid fish underwent a whole genome duplication ~100 million years ago, and presently about 50% of the retained gene duplicates have diverged regulation (3). In a recent study (5) we used a comparative phylogenetic approach to explore the impact of WGD on adaptive evolution of gene expression in the liver. Using the salmonids as a study system, we found that WGD boosted gene expression evolution, with the majority of duplicate pairs evolving lower expression in one of the duplicates across many tissues. These genes also showed significant enrichment of transposable element insertions in the promoter region. Only a very small fraction of duplicated genes (30 pairs) evolved liver-specific increase in expression in one copy, as expected under a scenario of adaptive evolution of novel gene functions following WGD. Interestingly, these candidates showed strong signatures of gains in transcription factor binding sites (TFBSs) for liver-specific transcription factors. Furthermore, TFBSs predicted to be bound by liver-specific TFs overlapped transposable elements more often in the evolved copy. These findings hint to a role of transposable elements in gene regulatory rewiring following WGD.

TE activity can have a large impact on evolution of TFBSs and *cis*-regulatory landscapes (6,7) and are sometimes co-opted by their hosts (7). In the short term, functional and active TEs can directly donate sequence motifs embedded in the TE that function as TFBSs in the host genome. At a longer time scale, TE insertions can accumulate secondary mutations that can result in *de novo* evolution of CREs. In Atlantic salmon, transposable elements make up a large fraction of the genome (about 51%), with the largest superfamily of Tc1-*Mariner* DNA-transposons experiencing an expansion coinciding with the WGD (3). This has further fuelled the hypothesis that transposable elements have played a key role in sequence- and regulatory divergence. Yet the potential of specific TEs as *cis*-regulatory elements (CREs) in salmonids is still unknown.

In this study we aim to test the hypothesis that transposable elements play a role in tissue-specific expression divergence among duplicate gene copies from the salmonid WGD. We do this using luciferase promoter reporter assays to test the *cis*-regulatory activity of transposable elements predicted to be bound by liver biased transcription factors identified in Gillard et al. (5).

## Results

### Tc1-*Mariners did not induce transcription*

Gillard et al. (5) found that the gene copies that had evolved increased expression levels in the liver were enriched for Tc1-*Mariner* insertions, which could act as binding sites for liver-active TFs, nearby or within the gene. To functionally evaluate if these Tc1-*Mariner* insertions could be responsible for evolution of increased liver transcription, we cloned four genomic copies into LUC reporter-assay vectors (Figure 1A). The four elements had the highest sequence similarity to Tc1-2, *Mariner*-22, and *Mariner*-10 families based on blast-searches to RepBase (<u>Supplementary file 1</u>) and verified through visual inspections of multiple sequence alignments.

**Figure 1.**
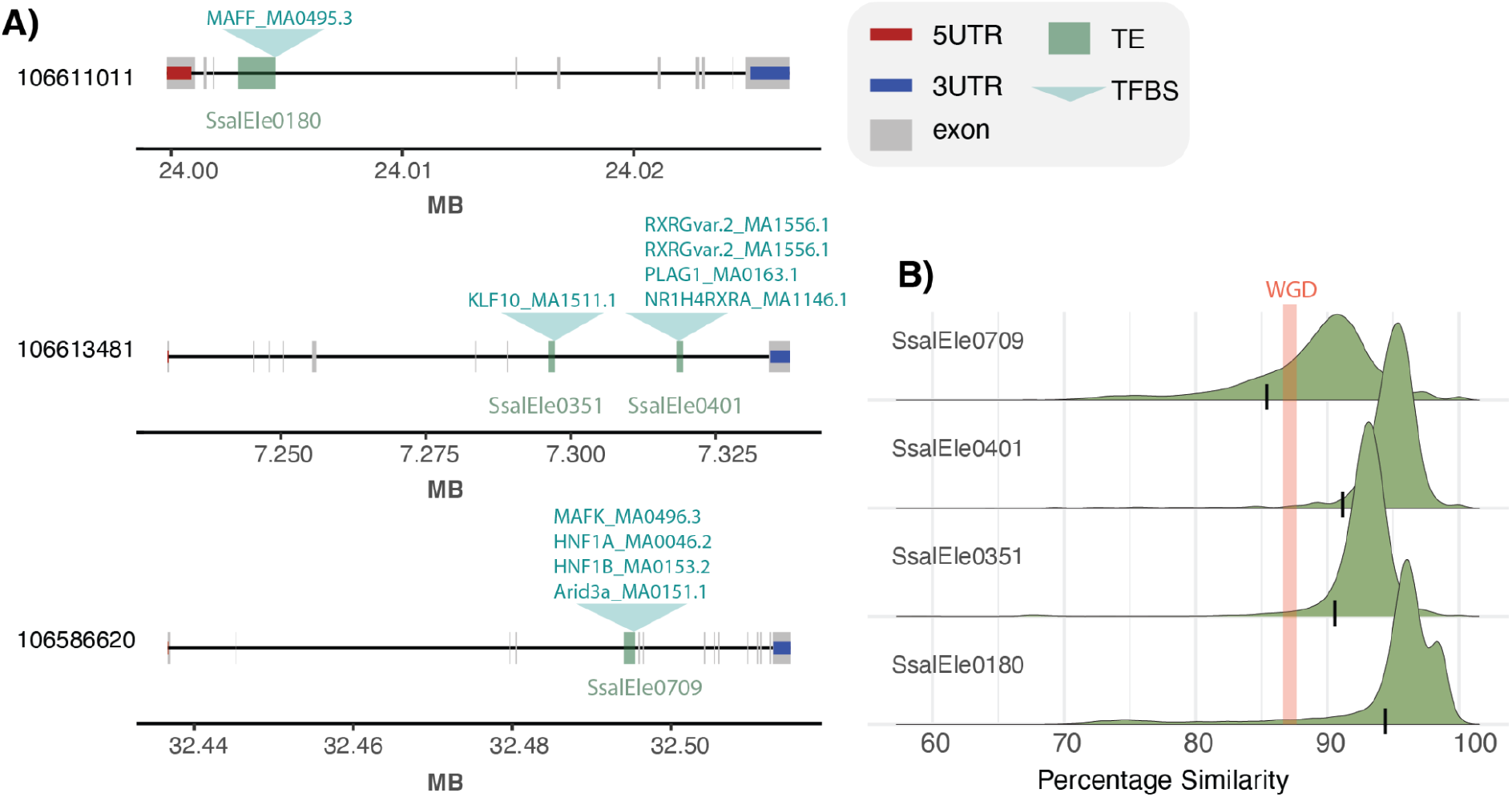
Overview of functionally validated Tc1-*Mariner* elements. A) Overview of element insertion relative to gene features (dark green) and predicted TFBS bound by liver-expressed TFs (light green above gene track). B) Within TE-family similarity distribution reflecting age of transposition activity. TE insertions in Figure 1A marked with a black line. Red line marks expected genomic similarity for duplicated genomic regions after WGD (3).

To assess the relative age of each TE subfamily activity and relative ages of the four respective element insertions, we compared the distribution of similarity between the genomic element copies and the subfamily consensus sequence (Figure 1B). The results showed that TE-subfamily level similarity peaks at >90% suggesting that these TEs were active after the WGD in salmonids (average similarity between duplicated genomic regions are 87% according to Lien et al. (3)). However, one of the cloned element copies (SsaEle0709) had a slightly lower similarity to the consensus sequence compared to what we expected for transposition events happening post-WGD (Figure 1B).

As a first experiment to evaluate the regulatory activity of the Tc1-*Mariner* superfamily elements we performed a LUC-reporter assay using the entire TE insertions (Figure 2A) as *cis*-regulatory elements. Whole TE sequences did not increase expression of the LUC reporter relative to the SV40 control (Figure 2A). Instead we observed a non-significant reduction in luciferase signals.

**Figure 2.**
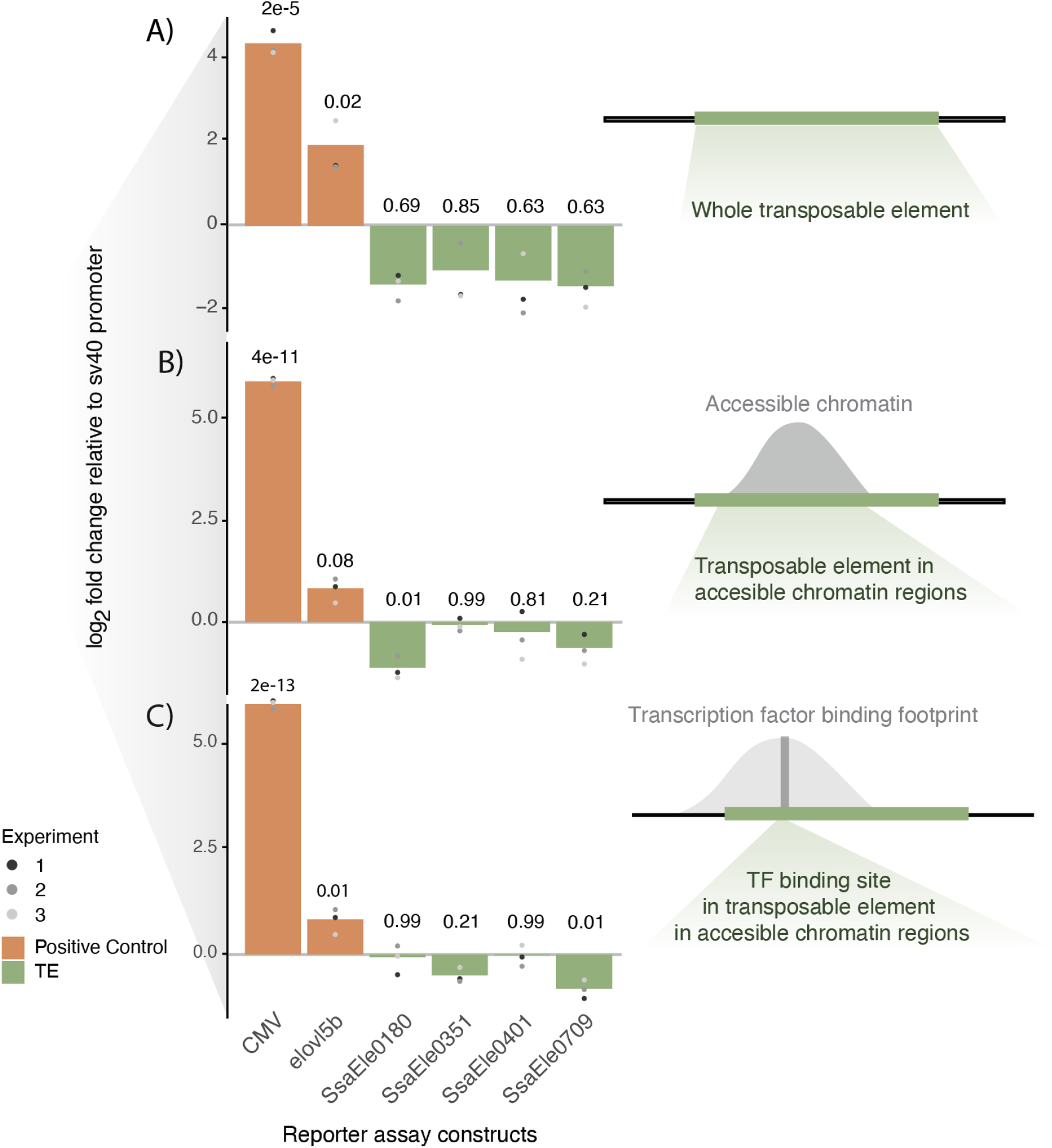
Dual luciferase reporter assays using Tc1-*Mariner* TE regions. Barplots showing log_2_ transformed mean fold change relative to sv40 empty vector across three experiments (A-C) with three replicates per experiment. (A) Reporter assays testing whole TEs. (B) Reporter assay testing accessible chromatin regions overlapping the TEs. (C) Reporter assay testing regions of the TEs in open chromatin and predicted to be bound by liver-specific TFs. Positive controls are the CMV enhancer and the ATAC-peak closest (upstream) to the transcription start site (TSS) of the liver expressed gene *elovl5b* (NC_059469.1 (ssa28: 27245480..27256837, Assembly ICSASG_v2, Gene ID: 100192340) from Atlantic salmon.

In their native states in salmon liver cells, only smaller regions of these Tc1-*Mariner* elements are in accessible chromatin regions and hence can act as potential binding sites for TFs (<u>Supplementary data file 2</u>). It is thus possible that by using whole TEs in the reporter constructs we introduce binding of transcriptional suppressors that can cloak effects of positive regulators. To explore this hypothesis we conducted two additional experiments using a nested design to exclude regions of the Tc1-*Mariner* elements likely not accessible to TFs in the native liver genome. We first tested regions of the TEs only within accessible chromatin peaks (i.e. ATAC-seq peaks), and finally, smaller regions overlapping these ATACseq peaks predicted to contain TFBSs bound by liver-expressed TFs (<u>Supplementary data file 2</u>, Figure 1 B-C). Similar to the whole TE experiment, sub-regions of the Tc1-*Mariner* elements did not induce transcription compared to our SV40 negative control (Figure 2B-C), and in two of the experiments significant repressive effects on luciferase signals were observed. In conclusion, we find no evidence supporting that the Tc1-*Mariner* elements tested here were involved in evolution of increased gene expression levels in the liver following the salmonid whole genome duplication.

### Non-*Mariner elements induce transcription*

Since the Tc1-*Mariner* elements associated with gene copies with increased expression levels did not induce transcription in reporter assays (Figure 2), we decided to test other putative TE-derived CREs associated with the same genes. Six TE-derived sequences situated 500bps upstream to 200bps downstream of the TSS and carrying TFBS predicted to be occupied by liver-expressed TF were synthesized, cloned into LUC-vectors, and tested for regulatory activity (Figure 3A). The existing annotation (3) classified these TE-CREs as one LINE1 superfamily element (SsalEle0377), one unknown DNA-transposon (SsalEle0849), and four TE-like sequences of unknown origin. Visual inspection of alignments (<u>Supplementary data files 3-12</u>) and blast searches to RepBase confirmed the annotation of the LINE1 superfamily element, the DTX-element annotation was considered low confidence, and the unknown TE-elements were confirmed to be too degenerated to be assigned to a superfamily.

**Figure 3.**
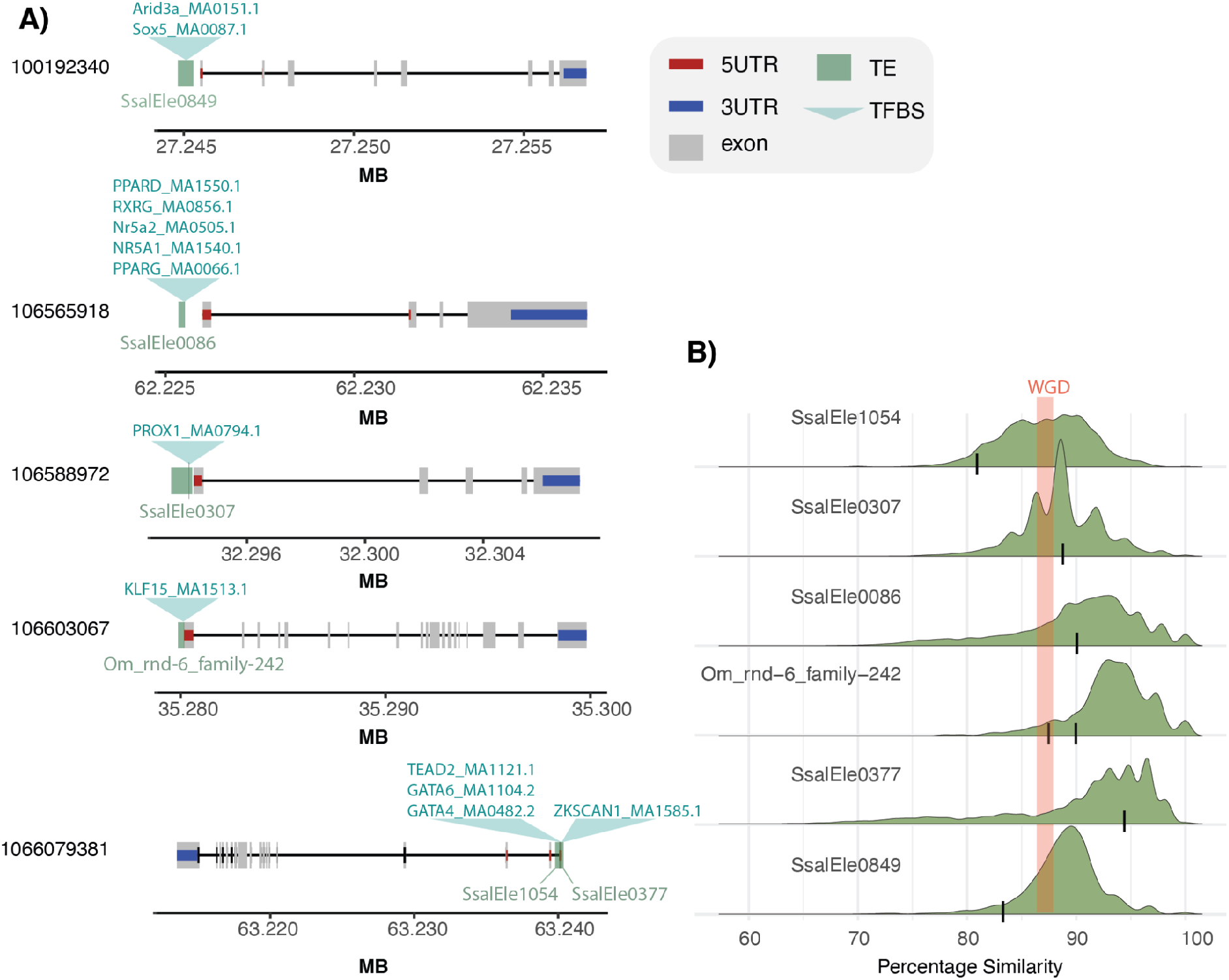
Overview of functionally validated non-*Mariner* elements. A) Overview of element insertion relative to gene features (dark green) and predicted TFBS bound by liver-expressed TFs (light green above gene track). B) Within TE-subfamily similarity distribution reflecting age of transposition activity. TE insertions in Figure 3A marked with a black line. Red line marks expected genomic similarity for duplicated genomic regions after WGD (Lien et al. 2016).

LUC-reporter assays showed significant positive effects on transcription for all constructs (p<0.0013) with mean fold change ranging between 2.78-35.37 (Figure 4). The SsaEle1054 had the smallest effect, while the LINE1-like SsalEle0377 had the largest effect on transcription. It is worth noting that we observed large variation in normalized luciferase signals between replicate experiments (Figure 4), however the rank order of effect sizes across experiments were relatively constant.

**Figure 4.**
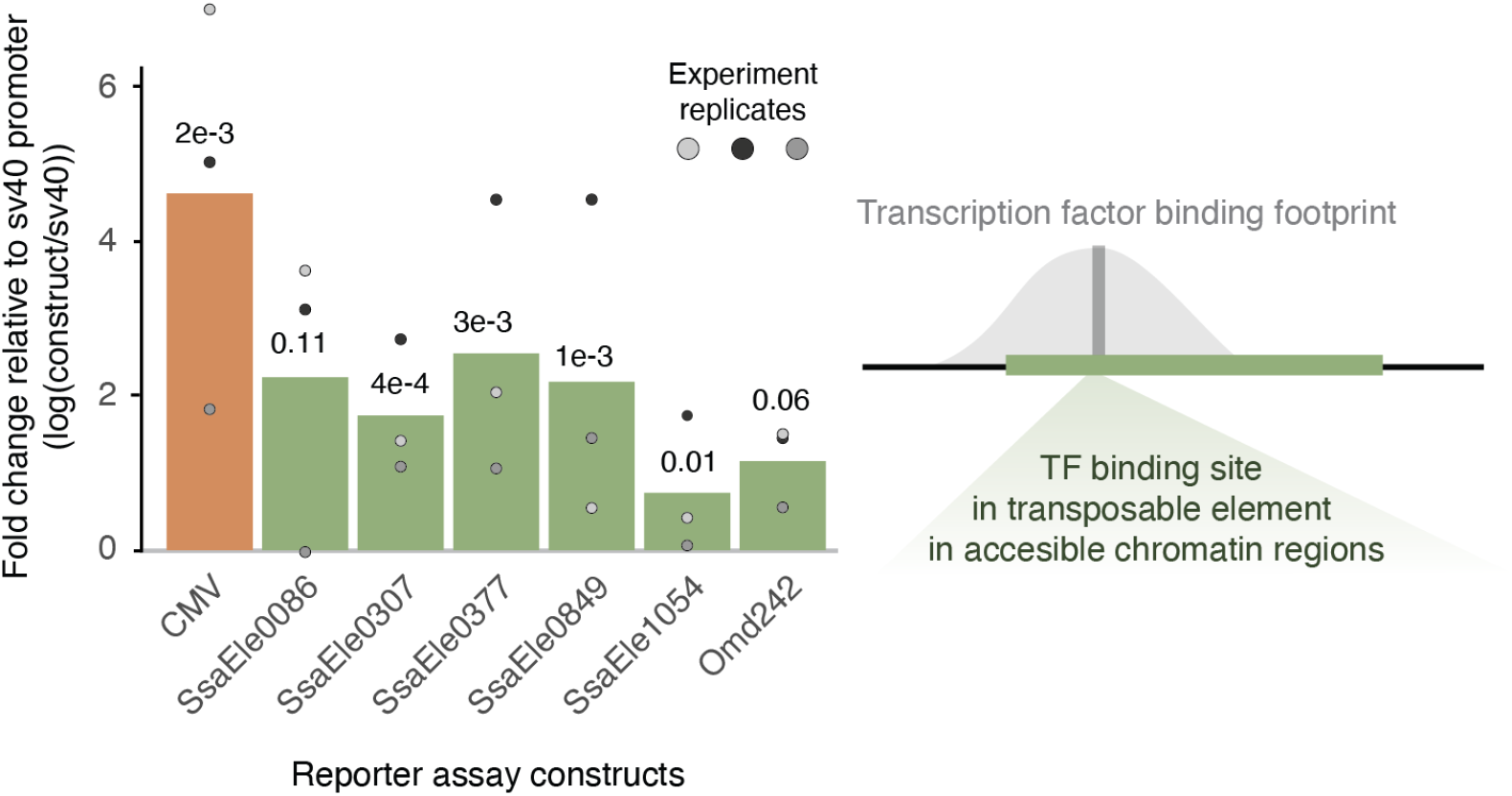
Dual luciferase assays on SINE and LINE candidates for TE-CREs. Barplots showing log_2_ transformed mean fold change relative to sv40 empty vector across three experiments with three replicates per experiment. Mean value for each experiment is indicated with a circle.

## Discussion

TEs are known to be important players in the evolution of gene regulation (7–10). In a recent paper we found that gene duplicates, where one copy evolved higher expression in the liver after a salmonid whole genome duplication, were enriched for insertions of Tc1-*Mariner* elements (5). Although Tc1-*Mariner* elements have been shown to function as promoters and can activate transcription (11), promoter reporter experiments in this study indicated no effect or very slight repressive effects on transcription for the four Tc1-*Mariner* superfamily elements tested (Figure 2 A-C). This is similar to what was found in a reporter assay experiment of primate TE-CREs in liver (Trizinno et al.2017), where only 16% (mostly LTRs and DNA transposons) induced transcription. The Tc1-*Mariners* elements tested here likely did not play an important role in evolution of liver specific divergence of gene duplicate expression as defined in Gillard et al. (5). It is worth noting, however, that the Tc1-*Mariner* elements tested in this study were derived from introns (Figure 1), and if TF-independent mechanisms such as intron-mediated enhancement (12,13) play a role in the gene duplicate divergence, our promoter-reporter assays would not be able to detect this.

The gene copies with increased liver expression also had other TEs with TFBSs predicted to bind liver-biased TFs in their promoters. Contrary to the Tc1-*Mariner* elements, these non-*Mariner* TE-CREs increased luciferase signals in our reporter assays (Figure 4). The TE-CRE with highest transcriptional induction, SsalEle0377, is a LINE1 element, which are known to function as CREs in other species (6). This TE-CRE only carried one TFBS predicted to be bound by a TF, a ZKSCAN1. Although we could not find literature supporting liver-specific function for this TFBS, the TF is highly expressed in the liver (according to GTEx) and associated with various roles in tumor development (14,15). The other TE-CREs with inductive effect on transcription all contained several TFBS motifs with known roles in liver cell gene regulation and function (Figure 4). The PPAR, Nr5a, and RXRa motifs in the SSalEle0086 element are known to play various roles in liver energy metabolism and induce transcription (16–18), while the GATA4, GATA6, and PROX1 motifs (in SsalEle1054 and SsalEle0307) are involved in liver cell fate specification and differentiation (19–21).

## Conclusion

TEs are suspected of playing an important role in rewiring gene regulation in the Atlantic salmon genome after a WGD event. Our results support that TE activity has contributed to liver specific gene duplicate divergence, but cast doubts about the importance of Tc1-mariners in gene regulatory rewiring following the Atlantic salmon WGD. Nevertheless, a more systematic genome-wide approach is needed to reach a general conclusion regarding all Tc1-mariner elements. Plasmid based reporter assays as used in this study also have clear limitations as they do not assay the CREs in a chromosomal context (22) and can not evaluate TF-independent gene regulatory mechanisms. Future studies could attempt to perform CRISPR based TE-copy specific knock-out to overcome these limitations.

## Materials and methods

### TE selection for reporter assay characterization

Gillard et al. (5) identified 30 Atlantic salmon gene duplicate pairs where one copy had evolved a liver-specific increase in expression following WGD. Among those genes several of the promoter regions were predicted to be bound by liver-active transcription factors at TFBSs within sequences annotated as TEs. Using this dataset as a starting point we first identified Tc1-mariner insertions that were candidates for being CREs and drive gene transcription in the liver. Fifty two bound TE-TFBSs located close to the transcription start sites (−500/+200bp’s) were identified. Of these, 5 Tc1-mariner elements were selected for reporter assays; SsalEle0180, SsalEle0256, SsalEle0351, SsalEle0401, and SsalEle0709. These were only present in the promoter of the gene duplicate copy with diverged expression.

When selecting non-mariner TEs for reporter assays we used the same list of putative TE-CREs. We required TE sequences to be >100bp and only present in the promoter region (−500/+200bp from transcription start site) of the gene copy with high liver expression. This resulted in 7 TE-CREs with a putative role in evolution of increased liver expression. One of these TEs contained a nested insertion of another TE fragment, both part of the OM_rnd-6 family-242, and these were tested together. TE-CRE filtering steps were carried out in R and the code can be found at https://gitlab.com/hansahls/te_reg_repository.

### TE characterisation

Manual curation of the selected TE insertions’ sequences were carried out using a pipeline described in Suh. et al (23): Blastn was first used to identify similar sequences in the genome. We selected the genomic regions of the 20 best hits and extracted these regions (+/- 2kb) as fasta sequences fasta format using the getfasta function in bedtools v2.18 (24), aligned sequences using MAFFT (25), and visually inspected the alignments. TEs were characterized according to terminal repeats, ORF compositions, known characteristic motifs and Repbase similarity searches (26).

To estimate a relative measure of the transposition activity for the TE subfamiles we used RepeatMasker (v. 4.1.0) (27) to compute the sequence similarity between TE insertions and their consensus TE sequence. We estimated the relative age (percent similarity at the base pair level) of the TE copies used in reporter assays by blasting the fasta for each TE insertion sequence to its consensus TE sequence.

### Phylogenetic analysis of Tc1-*Mariner* transposable elements

To get an overview of the relatedness and subclassification, we conducted a phylogenetic analysis of the Tc1-*Mariner* TEs (SsalEle0180, SsalEle0256, SsalEle0351, SsalEle0401, and SsalEle0709). DNA sequences of the TEs were extracted from the salmon genome sequence (NCBI: ICSASG v2) in the fasta format using the getfasta function in bedtools v2.18 (24). The TE sequences were then combined with all Tc1-*Mariner* consensus sequences from (3). TEs shorter than 100 bp were disregarded, and the sequences were then aligned using MAFFT v7.475 (25) with the parameters –adjustdirection and –auto. A phylogenetic tree was then estimated from the multiple sequence alignment using FastTree v2.1.11 (28). This tree was then visualized using the ggtree package (v2.2.4; (29)) in R. The code for this phylogenetic analysis can be found at https://gitlab.com/hansahls/te_reg_repository/-/blob/main/Phylogenetic_Analysis.Rmd

### Preparation of luciferase reporter constructs

To validate the transcriptional regulatory roles of the putative TE-CREs found to be associated with liver-specific duplicate gene expression, we performed luciferase reporter assays. The reporter vectors were constructed in two different ways, using PCR amplicons or oligo-synthesis. For five Tc1-*Mariner* elements we PCR-amplified whole TE-elements or sub-regions of the TEs from Atlantic salmon genomic DNA and cloned these into vectors for reporter assays. PCR were done using Platinum™ SuperFi ™ PCR Master Mix (Invitrogen) using primers that also bear 15 bp tail sequences homologous to SacI and XhoI restriction enzyme cloning sites within the pGL3-Promoter firefly luciferase vector (Promega, GenBank® Accession number U47298). PCR primers are listed in <u>Supplementary file 1</u>. PCR-amplified elements were gel-purified using the QIAquick® Gel Extraction Kit (Qiagen #28706) and cloned into SacI- and Xho-digested pGL3-Promoter vector using the Infusion®HD Cloning kit (Takara Bio #639650). Elements were cloned upstream of the sv40 promoter within the pGL3-Promoter vector. Vector design and map files can be found in <u>Supplementary data files 13-17</u>. Recombinant reporter constructs were isolated using the ZymoPURE™ Plasmid Miniprep Kit (Zymo Research, #D4210), following the manufacturer’s protocol with the following minor modifications. 50mL tubes were centrifuged at 4000rpm for 5 minutes, and the supernatant was discarded. 500μL of ZymoPure ™ P1 (Red) was used for pellet resuspension, and then transferred to a 3.0mL Lo-bind tube. 500μL of ZymoPURE P2 and ZymoPURE P3 was used instead of 250μL. 600μL of lysate was transferred to two tubes (total volume 1200μL). Final elution centrifugation was done at 11,000 x g for 3 minutes instead of 1 minute. After elution the DNA concentrations were measured using a Nanodrop 8000 spectrometer.

Another set of six potential TE-CREs were synthesized and cloned within SacI- and Xho-linearized pGL3-Promoter vector (outsourced to Genscript). All reporter constructs were confirmed by Sanger sequencing (LightRun Tube Sequencing Service, Eurofins).

### Transfection of salmon primary liver cells and Dual-Glo® luciferase assay

We reasoned that Atlantic salmon primary hepatocytes are ideal for validating the roles of the TEs in liver-specific regulation of duplicated genes owing to the lack of salmon liver continuous cell lines, and to the fact that they are of liver origin. Primary hepatocytes were isolated using a protocol optimized by Datsomor et al. (30). Cells were isolated from Atlantic salmon with an average weight and length of 30.3 grams and 310.9 cm, respectively. Approximately 1.0 - 1.5 x10^5^ primary hepatocytes were co-transfected per well in 24-well plates with 1.7 μg of each Tc1-*Mariner* reporter construct together with 0.3 μg of pGL4.75[*hRluc/*CMV] (Promega) which encodes Renilla luciferase whose activity is used as an internal standard for normalizing variations in cell number and transfection efficiency. For SINE and LINE reporter constructs, 1.5 μg of each reporter construct was co-transfected with 0.5 μg of pGL4.75[*hRluc/*CMV]. Transfection was performed using the Neon™ transfection system with an electroporation program optimized for primary hepatic cells (30): 1400 voltage, 20 ms pulse width and 2 pulses. The primary hepatocytes were cultured at 15 ^0^C under atmospheric conditions in L15 GlutaMAX™ medium (ThermoFisher) supplemented with 5% foetal bovine serum (without antibiotics). Medium was replaced with fresh L15 GlutaMAX™ supplemented with 5% foetal bovine serum and 1x penicillin-streptomycin 24 hrs post-transfection and the cells cultured for additional 24 hrs. To assess firefly luciferase activities per well, medium on cells was replaced with 100 μl each of Dulbecco’s Modified Eagle’s medium (Sigma) and Dual-Glo® luciferase reagent (Promega) and incubated for 30-45 min. Luminescence was read on Synergy H1 Hybrid multi-mode microplate reader (Bio Tek). Luminescence from Renilla luciferase activities was measured 10 min after adding 100 μl of Dual-Glo® Stop & Glo® reagent. Firefly luminescence was normalized to Renilla luciferase luminescence and presented as means of triplicates, unless stated otherwise.

### Statistical analyses of differences in luciferase expression

Initial processing of the luciferase assay results was done according to the manufacturer’s instructions (Dual-Glo® Luciferase Assay System, Promega, #E2920). In brief, normalization of the data was carried out by calculating the fold change between firefly-RLU and renilla-RLU for each well, and then calculating the mean of all replicates. This normalization step mitigated unwanted effects from differences in transfection efficiency and cell survival variability between wells. The signals from the untransfected wells acted as a control for well contamination and other experimental issues.

A statistical test for differences in LUC-expression was performed for all experiments. For the experiments with non-Tc1-*Mariner* TEs we performed the luciferase experiment three times, yielding large differences in absolute luciferase values. To account for this we therefore used a linear regression model, with each experiment as a cofactor to test whether the TEs have a significant effect on luciferase expression. For the Tc1-*Mariner* experiments, where we only performed one experiment with three replicates for three different constructs per TE (whole TE, ATAC-peak, TFBS-motif) we used an ANOVA test followed by a Post-Hoc Tukey Multiple comparison of means analysis. Scripts to reproduce statistical tests and visualizations can be found at https://gitlab.com/hansahls/master_thesis_hanna_sahlstrom/-/tree/master/R_code.

## Data and code availability

All data and code used to generate analyses and figures will be be available in a static figshare repository [DOI coming here] after peer review. Meanwhile all the data and code to reproduce results, figures and statistical analyses are available at https://gitlab.com/hansahls/te_reg_repository.

## Author contributions

SRS conceived the study. SRS, HS, and AKD designed the study. HS and AKD performed all of the lab work. HS, AKD, SRS, TRH and ØM performed data analyses. HS, AKD, TRH, and SRS drafted the manuscript. All authors revised the manuscript.

## Acknowledgments

This study was funded by the Norwegian Research Council through the projects Digisal (248792) and Rewired (274669).

## Supplementary files

*Supplementary file 1*: PCR primers for amplification of TE-associated DNA with putative CRE function. https://gitlab.com/hansahls/te_reg_repository/-/tree/main/Supplementary_Files

*Supplementary file 2*: Further information and sequence data on tested TEs https://gitlab.com/hansahls/te_reg_repository/-/tree/main/Supplementary_Files

*Supplementary files 3-12*: Sequence alignments used to confirm TE family annotation https://gitlab.com/hansahls/te_reg_repository/-/tree/main/Supplementary_Files/TE_Alignments

*Supplementary files 13-17: sequence* maps of vector constructs that were designed and used in luciferase assay analysis. https://gitlab.com/hansahls/te_reg_repository/-/tree/main/Supplementary_Files/Vector_maps

*Supplementary file 18*: Primers for Sanger sequencing of vector constructs to confirm accurate cloning of TE constructs into vector. https://gitlab.com/hansahls/te_reg_repository/-/tree/main/Supplementary_Files

